# Inferring latent temporal progression and regulatory networks from cross-sectional transcriptomic data of cancer samples

**DOI:** 10.1101/2020.10.07.329417

**Authors:** Xiaoqiang Sun, Ji Zhang, Qing Nie

## Abstract

Unraveling molecular regulatory networks underlying disease progression is critically important for understanding disease mechanisms and identifying drug targets. The existing methods for inferring gene regulatory networks (GRNs) rely mainly on time-course gene expression data. However, most available omics data from cross-sectional studies of cancer patients often lack sufficient temporal information, leading to a key challenge for GRN inference. Through quantifying the latent progression using random walks-based manifold distance, we propose a latent-temporal progression-based Bayesian method, PROB, for inferring GRNs from the cross-sectional transcriptomic data of tumor samples. The robustness of PROB to the measurement variabilities in the data is mathematically proved and numerically verified. Performance evaluation on real data indicates that PROB outperforms other methods in both pseudotime inference and GRN inference. Applications to bladder cancer and breast cancer demonstrate that our method is effective to identify key regulators of cancer progression or drug targets. The identified ACSS1 is experimentally validated to promote epithelial-to-mesenchymal transition of bladder cancer cells, and the predicted FOXM1-targets interactions are verified and are predictive of relapse in breast cancer. Our study suggests new effective ways to clinical transcriptomic data modeling for characterizing cancer progression and facilitates the translation of regulatory network-based approaches into precision medicine.

**Author summary:** Reconstructing gene regulatory network (GRN) is an essential question in systems biology. The lack of temporal information in sample-based transcriptomic data leads to a major challenge for inferring GRN and its translation to precision medicine. To address the above challenge, we propose to decode the latent temporal information underlying cancer progression via ordering patient samples based on transcriptomic similarity, and design a latent-temporal progression-based Bayesian method to infer GRNs from sample-based transcriptomic data of cancer patients. The advantages of our method include its capability to infer causal GRNs (with directed and signed edges) and its robustness to the measurement variability in the data. Performance evaluation using both simulated data and real data demonstrate that our method outperforms other existing methods in both pseudotime inference and GRN inference. Our method is then applied to reconstruct EMT regulatory networks in bladder cancer and to identify key regulators underlying progression of breast cancer. Importantly, the predicted key regulators/interactions are experimentally validated. Our study suggests that inferring dynamic progression trajectory from static expression data of tumor samples helps to uncover regulatory mechanisms underlying cancer progression and to discovery key regulators which may be used as candidate drug targets.

## Introduction

Inferring gene regulatory networks (GRNs) from molecular profiling of large-scale patient samples is of significance to identifying master regulators in disease at systems level [1, 2]. Detecting the causal relationships between genes from biomedical big data, such as clinical omics data, has recently emerged as an appealing yet unresolved task, particularly for clinical purposes (e.g., diagnosis, prognosis and treatment) in the era of precision medicine [3].

Many methods have been developed for inferring GRNs from gene expression data [4]. The GRN inference methods can be grouped into at least four categories: Boolean network methods [5], ordinary differential equation (ODE) model-based methods [6], Bayesian network methods [7] and tree-based ensemble learning methods [8]. These methods mainly rely on two types of gene expression data, i.e., gene perturbation experiments [9, 10] or time-course gene expression data [11]. Temporal changes in expressions, resulting from the interactions between genes, could potentially imply causal regulations. Meanwhile, a wealth of time-course transcriptomic data has been generated from the laboratory experiments. So temporal type of expression data is one of the most common assumptions based on which many GRN inference methods were designed [12].

However, the transcriptomic data of tumor samples often lack explicit temporal information [13]. In fact, large samples of time-course data are rarely available in clinical situations, at least for now, since longitudinal surveys are often challenging to conduct. In contrast, cross-sectional studies (i.e., a snapshot of a particular group of people at a given point in time) based on high-throughput molecular omics data are more prevalent due to their relative feasibility. As such, for cross-sectional transcriptomic data at population-scale, most of the current methods, such as Pearson correlation coefficient (PCC)-based methods [14], mutual information [15], regression methods [16] and machine learning methods [17], can only infer co-expressions or associations between genes. Moreover, although some correlation network-based methods have been used to identify disease-associated genes [18], it’s hard to tell the causal drivers or regulatory roadmap underlying phenotypic abnormality in the absence of regulatory network information [19]. Therefore, the lack of temporal information in clinical transcriptomic data leads to a key challenge for inferring directed GRN and its translation to systems medicine.

Decoding temporal information that traces the underlying biological process from the cross-sectional data is intriguing and enlightening to address the above challenge. The sample similarity-based approach has shown great promise in recovering evolutionary dynamics in evolution and genetics studies [20], for instance, phylogenetic trees based on microarray data [21] and genetic linkage maps based on genetic markers [22]. To this end, we propose that the latent temporal order of cancer progression status (i.e., latent-temporal progression) could be estimated from the cross-sectional data based on transcriptomic similarity between patient samples. Leveraging the latent-temporal ordering, we could represent the GRN as a nonlinear dynamical system. What’s more, however, considering the technical variability or measurement error in the RNA-sequencing or microarray data (e.g., variations in sample preparation, sequencing depth and measurement noise and bias) [23, 24], it’s indispensably important to guarantee the robustness of the GRN inference.

In this study, we present PROB, a latent-temporal progression-based Bayesian method of GRN inference designed for population-scale transcriptomic data. To estimate the temporal order of cancer progression from the cross-sectional transcriptomic data, we develop a staging information-guided random walk approach to efficiently measure manifold distance between patients in a large cohort. In this way, the cross-sectional data could be reordered to be analogous to time-course data. This transformation enables us to formulate the GRN inference as an inverse problem of progression-dependent dynamic model of gene interactions, which is solved using a Bayesian method. The robustness of the estimates of regulatory coefficients is justified through mathematical analysis and simulations. Furthermore, applications to real data not only demonstrate better performance of PROB than other existing methods but also show good capacity of PROB in identifying key regulators of cancer progression or potential drug targets. The identified ACSS1 in bladder cancer and predicted FOXM1-targets interactions in breast cancer are both validated. In addition, we also discuss potential clinical applications of our method.

## Materials and Methods

### Latent-temporal progression-based Bayesian (PROB) method to infer GRN

#### Overview of PROB

PROB consists of two major components. First, to infer the latent temporal information of cancer progression from the cross-sectional data, a graph-based random walk approach was developed to quantitatively order patient samples (**Figure 1a-b**). To this end, we defined a manifold distance between patients by analytically summing the transition probabilities over all random walk lengths to quantify temporal progression and the root was automatically identified with the aid of staging information. The quantitative reordering of the samples led to the recovery of the temporal dynamics of gene expression (**Figure 1c**). Second, a progression-dependent dynamic model was proposed to mechanistically describe the gene regulation dynamics during the above estimated temporal progression. To ultimately infer the GRN, the inverse problem in terms of parameter estimation of the dynamic model was transformed to a linear regression model which was solved using a Bayesian Lasso method (**Figure 1d**). Compared to the existing correlational network methods, PROB can infer causal GRNs with directed and signed edges from cross-sectional transcriptomic data.

**Figure 1.**
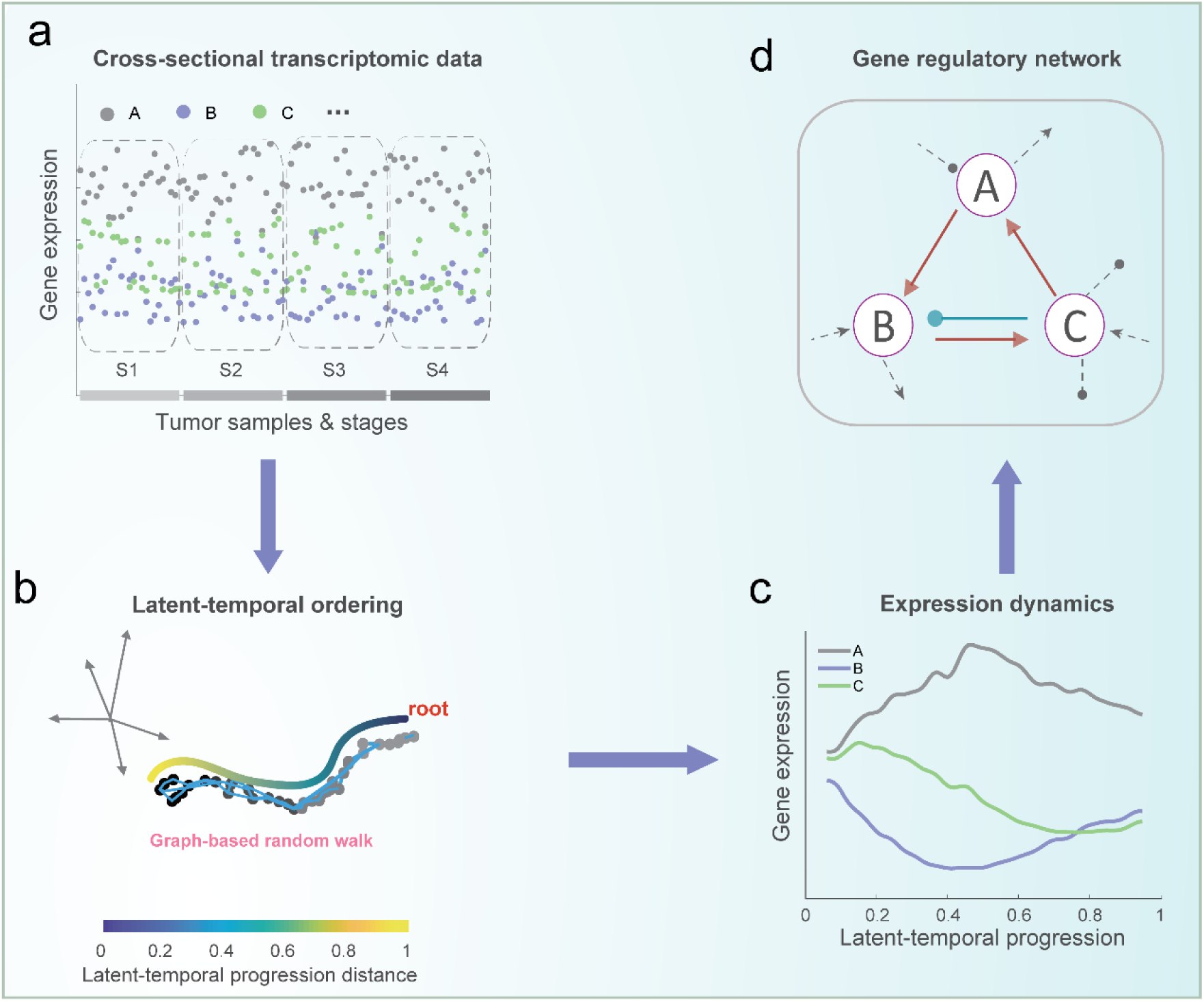
Illustration of the PROB framework for inferring the causal gene regulatory network from cross-sectional transcriptomic data. (**a**) Illustration of cross-sectional transcriptomic data, taking three genes (i.e., A, B, and C) as an example. Each sample was labeled with staging information (e.g., S1, S2, S3, and S4). (**b**) Similarity graph-based random walk approach for cancer progression inference. A scale-free temporal progression distance (TPD) is defined by analytically summing the transition probability between patients over all random walk lengths. Patients are thus ordered according to the TPD with respect to the root identified with the aid of staging information. (**c**) The expression dynamics of each gene according to the latent-temporal progression are then recovered. (**d**) A Bayesian Lasso method is developed to infer the causal GRN based on the temporal data of gene expression. Besides edge directions, PROB can also infer signs of the interactions (activation or inhibition), compared to the existing correlational network methods.

#### Temporal progression inference for cancer samples

We employ a similarity graph-based random walk approach to order patients along with the progression and to estimate the progression score for each patient, given the hypothesis that the similarity between patients can be measured by the patients’ gene expression profiles and pathology information. We first define a Gaussian similarity function for two patients, *x* and *y*, as

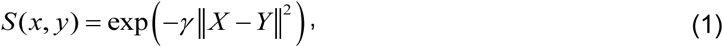

where *X* and *Y* are vectors used to represent the transcriptomic profiles of the respective patients and ‖*X* − *Y*‖ is the *L*^2^ norm of *X* − *Y*. The parameter *γ* is determined as

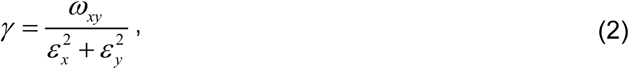

where *ω*_*xy*_ is a weight coefficient given by the available pathology information such as stage, which is defined in this study as *ω*_*xy*_=1 + |*G*_*x*_− *G*_*y*_|,with *G*_*x*_ and *G*_*y*_ representing staging information (taking values of, for instance, 1, 2, 3, or 4) of the two patients *x* and *y*, respectively. The parameter *ε*_*x*_ is adaptive for each patient *x* and is set as the patient’s distance to the *κ-*th nearest neighbor. *S* can be viewed as a stage-weighted and locally scaled Gaussian kernel.

Based on the above Gaussian kernel and normalization procedure (**Text S1**), we derive a transition probability matrix *P*, with element *P*_*xy*_ representing the probability of transitioning from patient *x* to patient *y* (or from *y* to *x*). We then measure the transitions on all length scales of random walks between patients. The accumulated transition probability (*Qxy*) of visiting *y* from *x* over random walk paths of all lengths is analytically calculated as

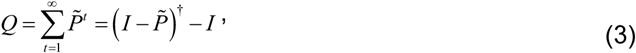

Where 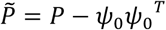, and *ψ*_0_ is the first eigenvector of *P* (corresponding to eigenvalue 1). Since *ψ*_0_ is associated with the steady state and contains no dynamic information [25], we subtract the stationary component *ψ*_0_*ψ*_0_^*T*^ from *P*, resulting in 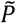. In this way, all the eigenvalues of 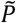 are smaller than 1; hence, the above sum of infinite series is convergent. 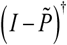 is the generalized inverse (or Moore-Penrose inverse) of 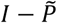 [26].

We use *Q*(*x*,·) to represent the accumulated transition probability of visiting all points from *x*. Thus, *Q*(*x*,·) is a row in *Q* and can be viewed as a feature representation for patient *x*. Therefore, we define a temporal progression distance (TPD) between two patients as

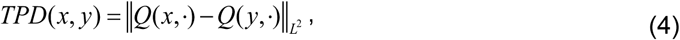

where ‖·‖ stands for the *L*^2^ norm. We remark that TPD is a scale-free manifold distance and is computationally efficient due to the closed form expression of *Q*.

Given a patient *x*, the progression score with respect to the root *x*_*0*_ is *s*=*LPD*(*x*_0_,*x*). Therefore, it is critical to determine the root sample in a large cohort for ordering the patients. We fulfill this task with the aid of the staging information of the tumor samples: among all patients, the root should have the largest TPD to a patient with maximal grade (e.g., grade 4). That is, the root *x*_0_ can be identified according to the following formula:

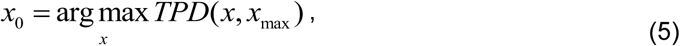

where *x*_*max*_ is a randomly selected patient from the maximal grade subpopulation.

We remark that the incorporation of staging information into the Gaussian kernel and root identification could significantly improve the accuracy of temporal progression inference (see **Figure S1** and Discussion section).

#### Dynamical systems modeling and parameter estimation

Based on the mass action kinetics [27], the temporal regulation of gene expressions can be modeled using the following dynamical system,

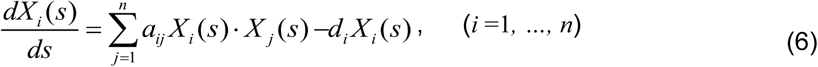

where *X*_*i*_(*s*) represents the expression level of gene *i* (*i* =1,*…,n*) in cancer with progression status *s. a*_*ij*_ is the regulatory coefficient from gene *j* to gene *i* (*j*≠*i*), and *d*_*i*_ is the self-degradation rate of gene *i*. The details of model assumption and derivation are provided in **Text S1** (“Progression-dependent dynamic modeling of the GRN” subsection).

Take *m*+1 points *S*_*i*_ = *s* (*r*_*i*_)(*i* = 0,1,, *m*) from the smoothed pseudotemporal progression trajectory *s* (*r*), where *r*_*i*_ = *i/ m*. We approximate 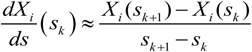 and denote 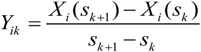, where *s*_*k* +1_ − *s*_*k*_ is sufficiently small (since *m* could be chosen large enough). Therefore, the above continuous model (i.e., Equation (4)) can be discretized and rewritten as

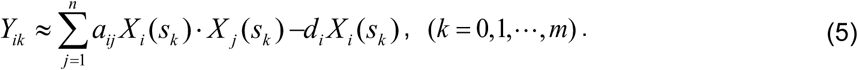

The above model is then transformed into a linear regression model and a scalable Bayesian Lasso method is adapted to estimate the posterior distributions of parameter values of *a*_*ij*_ and *d*_*i*_ for GRN reconstruction. See details in **Text S1** (“Parameter estimation using Bayesian Lasso method” subsection).

#### Mathematical analysis

Considering the technical variability or measurement error in the transcriptomic data [23, 24], it is important to examine the robustness of the method with respect to the perturbation in latent-temporal progression. To this end, we present the following theorem.

##### Theorem 1.

Assume there are two trajectories of latent-temporal progression *s*(*r*) and 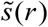 with the same root, *r* ∈ *I* = [0,1]. Define 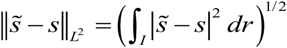. If (*X*_*i*_ (*s*), *a*_*ij*_)and 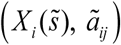 both satisfy the equations of progression-dependent dynamic model, i.e.,

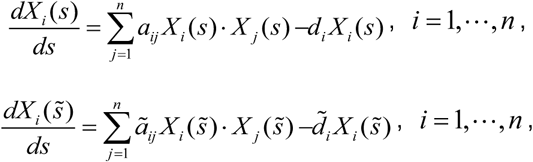

then we have

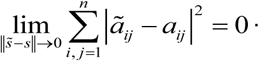

The proof of the above theorem is provided in **Text S2**.

Based on the spectral graph theory [28, 29], the above manifold distance (TPD) is noise-resistant, so the variation in the progression trajectory (i.e., 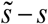) should be small given moderate perturbations (as illustrated below). Consequently, **Theorem 1** then implies that the corresponding estimates of 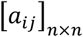 should vary minimally. Therefore, the above theorem theoretically guarantees the consistency and robustness of the estimates of the regulatory coefficients. In addition, the Bayesian Lasso method adopted by PROB further ensures a robust implementation of GRN inference.

#### Computational algorithm

The algorithm to infer progression trajectory and GRN is presented below. The implementation of PROB is described in **Text S3**.

##### Algorithm 1. pseudo-code of PROB

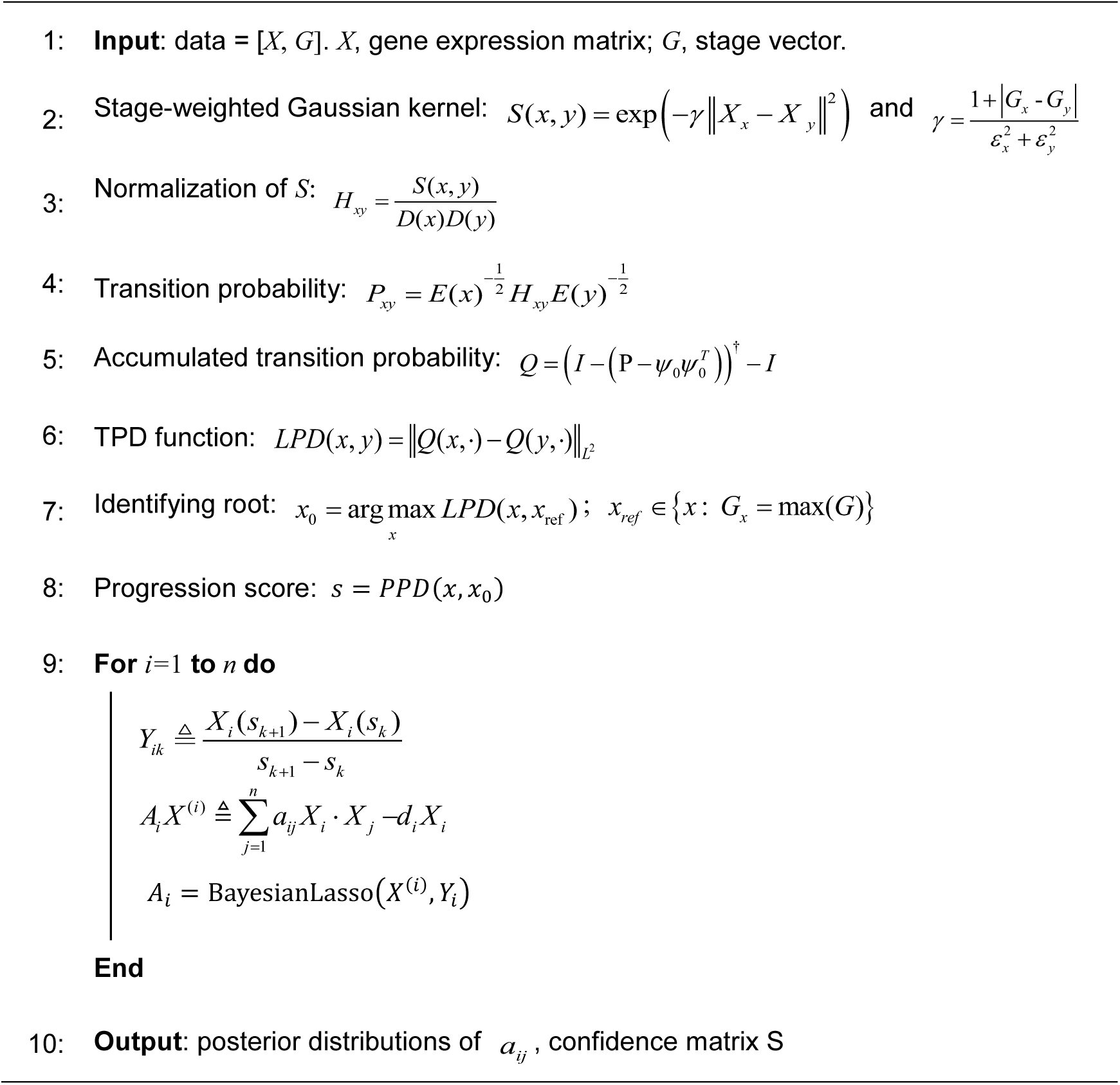

### Benchmarking PROB with alternative methods of GRN inference

For tumor sample-based gene expression data, several methods have been developed to infer gene networks. Pearson correlation (PCOR) is often used to quantify gene coexpression. Mutual information (MI) measures non-linear dependency between genes and thus provides a natural generalization of the correlation. MI-based methods for GRN inference include ARACNe [30], CLR [31], and MRNET [32]. Another commonly used method for GRN inference based on gene expression data is multiple linear regression LASSO method [16], which assumes sparse network structure and is feasible for high-dimensional data. Ensemble learning methods, such as GENIE3 (a tree-based ensemble learning method [17]), have been developed to infer gene regulatory relationships by viewing GRN reconstruction as a classification problem. In addition, we also included some GRN inference methods recently developed for scRNA-seq data into benchmarking analysis, since scRNA-seq data is also cross-sectional type. Such methods include SCODE [33] that uses ordinary differential equations model and LEAP [34] that constructs gene co-expression networks by using the time delay involved in the estimated pseudotime of the cells. SINCERITIES [35] is designed for time-stamped scRNA-seq data but requires at least 5 time points, so it is not applicable for the following benchmarking dataset as well as the tumor sample-based transcriptomic data.

In this study, we compared the accuracy of PROB with that of PCOR, ARACNe, CLR, MRNET, Lasso, GENIE3, SCODE and LEAP based on a real scRNA-seq data of dendritic cells (DCs) (GSE41265 [36]). The cells were stimulated with LPS and sequenced at 1, 2, 4, and 6h after stimulation. Only wild type cells (*n*=479) without Stat1 and Ifnar1 knockout were chosen for analysis. We choose this DC dataset for benchmarking because regulatory potential between 23 TFs in the DCs has been determined via a high-throughput Chromatin ImmunoPrecipitation (HT-ChIP) method [37]. The AUC of ROC was used to assess and compare the prediction accuracies of the above methods.

In addition, we collected a set of known regulators and targets [38] to test whether PROB could correctly distinguish outgoing regulations of different genes. To this end, we defined an outgoing causality score (OCS) for gene *i* in cell *k* as follows: 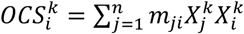, where *m*_*ji*_ is the mean of the posterior distributions of *a*_*ji*_, 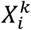 is the expression level of gene *i* in cell *k*. We then compared the distributions of OCS values of 6 regulators and that of 28 targets using the above DC dataset. The Wilcoxon rank-sum test (one-tailed) *p* value was calculated to assess statistical significance.

### Application to a dataset of bladder cancer

We applied PROB to a dataset of bladder cancer patients that includes 84 cases of conventional UCs and 28 cases of SARCs which were profiled by Illumina HumanHT-12 DASL Expression BeadChips (GSE128192 [39]). The temporal progression inference was performed to quantitatively order samples based on the whole gene expression profile with UC samples and SARC samples labeled by 1 and 2 respectively. To reconstruct epithelial-to-mesenchymal transition (EMT) regulatory networks, we collected 44 representative genes of TGFB1 pathway, RhoA pathway, p53 pathway, p63 pathway and EMT transcriptional regulators (**Table S1**) [39]. The UC network and SARC network were reconstructed based on the ordered expression data of the above 44 genes in UC samples and SARC samples respectively. The UC-specific network and SARC-specific network were then constructed by extracting edges that were unique to UC network and SARC network respectively. The out-degree values for each node in the two networks were calculated to poetize key regulator genes.

### Application to a dataset of breast cancer

We applied PROB to a microarray dataset of breast cancer (GSE7390 [40]) to identify key regulator genes with prognostic role in cancer progression. We identified the hub gene in the GRN based on an eigenvector centrality measure according to singular value decomposition method [41]. Denote the mean of the posterior distributions of *a*_*ij*_ as *m*_*ij*_, and 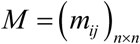. We subject *M* to singular value decomposition. We calculated the principal eigenvector of *MM*^*T*^ and denoted it *H=*(*h*_*1*_, *h*_*2*_, …, *h*_*n*_). The hub score of node *i* was defined as *h*_*i*_. The gene with greatest hub score was identified as a hub gene for further analysis and validation.

### Validation of the role of ACSS1 in bladder cancer

#### Antibodies and reagents

Anti-β-actin Mouse mAb (1:1000, 0101ES10, Yeasen), anti-E-Cadherin Mouse mAb (1:1000, #14472, CST), anti-ACSS1 Rabbit mAb (1:1000, 17138-1-AP, Proteintech), Goat Anti-Rabbit IgG (H+L) (1:10000, 33101ES60, Yeasen), Goat Anti-Mouse IgG (H+L) (1:10000, 33201ES60, Yeasen), Anti-Rabbit IgG-HRP kit (SV0002, Boster).

#### Over-expression plasmids and siRNA transfection

5637 cells were placed in 24 wells plate and transfected with the lentiviral vectors pTSB-CMV-puro and SiRNA against ACSS1 reaching 70%-80% confluence using Lipofectamine 2000 (Thermo Scientific) according to the manufacturer instructions. The SiRNA sequence used in this study are listed in Supplementary **Table S2**.

#### RNA extraction and qPCR

Total RNA was extracted by HiPure Total RNA Mini Kit (R4111-03, Magen) and the concentration was detected by ultramicrospectrophotometer (NanoDrop 2000, Thermo Fisher Scientific). RT-PCR was performed using PrimeScript RT Master Mix (DRR036A, TakaRa) and qPCR was performed by qPCR SYBR Green Master Mix (11198ES03, Yeasen) in Real-time quantitative PCR instrument (Q1000+, Long Gene). All the relative mRNA expression was normalized to GAPDH. The qRT-PCR primer sequence used in this study are listed in Supplementary **Table S3**.

#### Western blotting

Total protein was extracted by RIPA lysis buffer (JC-PL001, Genshare) with PMSF (1:100, 20104ES03, Yeasen). Standard western blot protocols were adopted. The band intensity of western blots was detected by BLT GelView 6000M. All the relative protein expression was normalized to β-actin.

#### Immunohistochemistry

All the tumor tissues were received from the operative resection of patients. The patients/participants provided their written informed consent to participate in this study. The studies involving human participants were reviewed and approved by the Ethics Committee of Sun Yat-sen University Cancer Center (approval no. GZR2018-131). The immunohistochemical analysis of the two biomarkers including ACSS1 and E-Cadherin was performed. All the pathological sections were produced, scan and analyzed by Leica Biosystems.

### Validation of the FOXM1 sub-network predictions

We validated the regulation of FOXM1 (a hub gene, see Results section) on the predicted targeted genes using multiple sets of gene expression data and ChIP-seq data that are publicly available.

To validate the expression changes of the predicted targeted genes following FOXM1 perturbation, we analyzed microarray gene expression data in MCF-7 cells that were treated with DMSO (control) or Thiostrepton (FOXM1 inhibitor) for 6 hours (GSE40762 [42]). The differential expression of the above 8 genes between control condition and FOXM1 inhibition condition was examined to test whether they were down-regulated after FOXM1 inhibition. The statistical significance was assessed using Wilcoxon rank sum test (one-tailed) p values.

To test whether FOXM1 binds to some of the predicted targeted genes, we used ChIP-seq data in both MCF-7 cell line (ER+) and MDA-MB-231 cell line (ER-) (GSE40762 [42]) to analyze binding of FOXM1. A standard procedure of the ChIP-seq analysis was performed for peak calling (**Text S7**).

## Results

### Testing PROB with a synthetic dataset

To illustrate the function of PROB, we generated a set of synthetic cross-sectional expression data (**Text S4**). For visualization purpose, we considered 6 genes in 100 cancer patients (**Figure S2a-b**). We first used PROB to infer temporal progression from the randomized sample-based data. The inferred latent-temporal progression was compared against the true progression (**Figure S2c)**, showing that PROB faithfully recovered the true ordering of the samples (Spearman’s rho=0.9991). The gene expression dynamics along with latent-temporal progression (**Figure S2d**) exhibited a very similar profile to the original data (**Figure S2a**). Based on the inferred temporal data, PROB inferred a GRN using the Bayesian Lasso method (**Figure S2e**). The posterior distributions of the regulatory parameters against their true values show that the estimation was rather reliable (**Figure S3**). An edge was determined by examining whether the 95% credible interval (CI) of the parameter estimates did not contain zero (**Text S4**). **Figure S2f** further demonstrates the accuracy of PROB in terms of GRN inference. The area under curve (AUC) of receiver operating characteristic (ROC) could be calculated for the inferred network compared with the ground-truth network based on the *k*% CI that contained zero or not.

To verify the robustness of PROB to the measurement variability, we further tested PROB for datasets at different levels of variabilities (**Figure 2**). The gene expressions were randomly perturbed by using multiplicative Gaussian noises to simulate different levels of measurement variabilities in the data, resulting in a series of coefficient of variations (CVs) (i.e., 0%, 5%, 10% and 15% respectively) (**Figure 2a**). The IDs of the samples were randomized to mimic sample-based snapshots of gene expression data, but the staging information was retained for each patient. PROB was applied to infer the GRN for each dataset. The accuracy of GRN inference was evaluated using the AUC of the ROC, showing that PROB could strongly reduce bias in gene expression measurements (**Figure 2b-c**) and robustly reconstructed the GRNs (**Figure 2d**).

**Figure 2.**
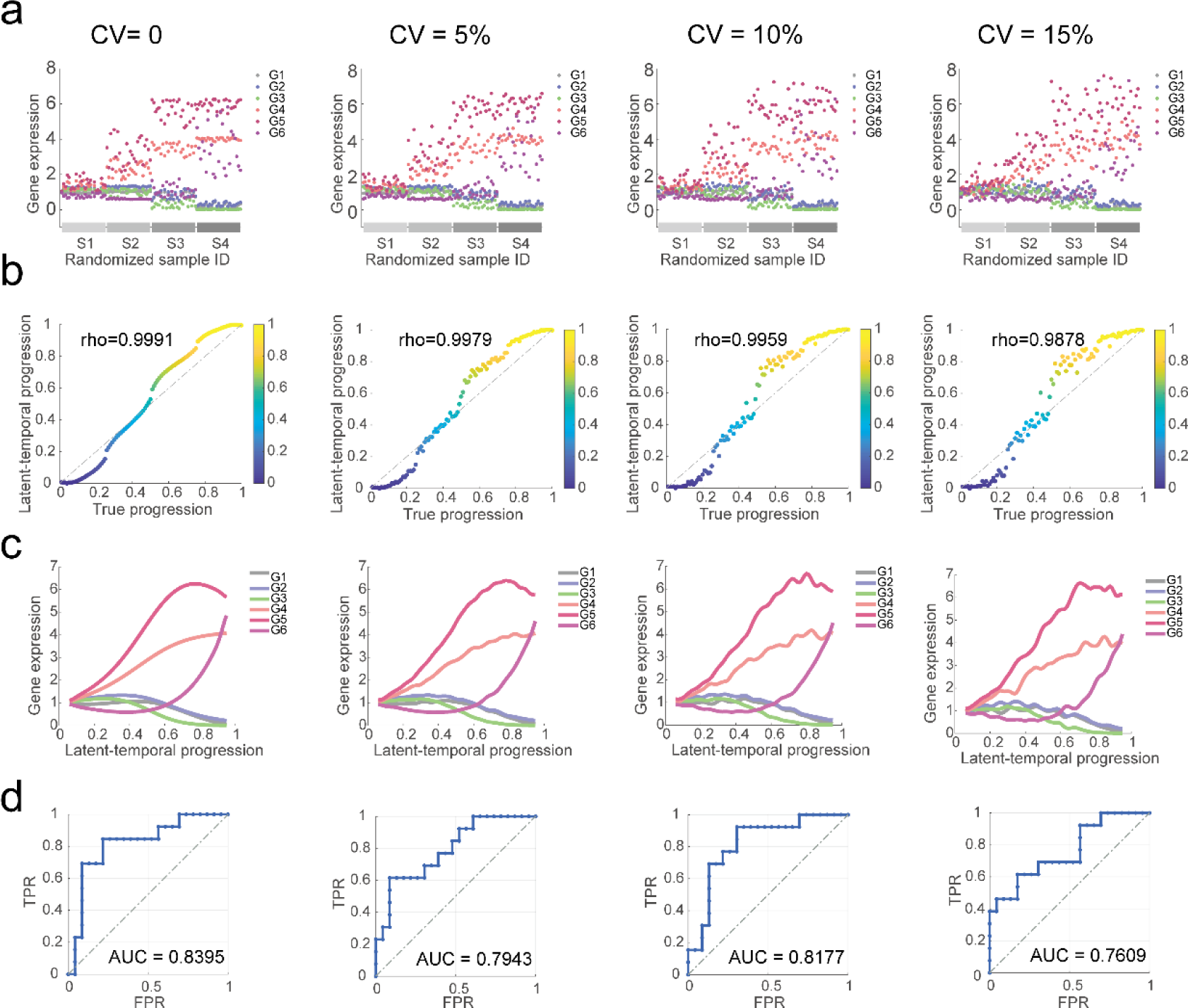
Demonstrating robustness of PROB using synthetic datasets at different levels of variabilities. A set of expression data for 6 genes in 100 cancer patients was simulated. Different levels of technical variabilities (with coefficient of variations (CVs) = 0%, 5%, 10% and 15% respectively) were introduced into the progression-dependent gene expression dynamics. (**a**) Simulated cross-sectional gene expression data. The sample IDs of the synthetic data were randomized and the staging information was retained. (**b**) Comparison of the inferred latent-temporal progression with the true progression in the synthetic dataset, evaluated using Spearman’s rank correlation coefficient (rho). (**c**) Recovered gene expression dynamics according to inferred progression trajectory. (**d**) Accuracy of the GRN inference evaluated using the areas under curve (AUCs) of the ROCs.

Additional evaluation metrics were employed to verify the robustness of PROB against a series of variations in the data (with CVs ranging from 0% to 30%). The root mean square error (RMSE) and Spearman correlation coefficients were used to evaluate the accuracy of the temporal progression inference (**Figure S4a-b**). The accuracy, positive predictive value (PPV) and Matthews correlation coefficient (MCC) were used to evaluate the robustness of the GRN reconstruction (**Figure S4c-f**). The findings are consistent with the above results (**Figure 2**).

### Benchmarking PROB with other existing methods

We used a set of single cell RNA-seq (scRNA-seq) data (GSE48968 [36]) for benchmarking of GRN inference methods since our method can be naturally applied to stage-stamped or time-course scRNA-seq data and the ground-truth of the GRN is available in this case as described in the Methods section. The LPS-stimulated dendritic cells (DCs) were sequenced at 1, 2, 4, and 6h after stimulation. The capture time in the data was treated as an analogy to ‘staging’ information when using PROB. The estimated latent-temporal progression recapitulated the physical progression of cells with a high correlation to the capture times (R^2^ =0.851) (**Figure 3a**). We compared PROB with other pseudotime inference methods (Slice, Slicer, PhenoPath, Wishbone, PAGA, Monocole2, DPT, Tscan). PROB estimation achieved a highest correlation with the original physical capture times among all methods tested, evaluated using both Kendall Tau rank correlation coefficient (**Figure 3b**) and coefficient of determination R^2^ (**Figure S5**).

**Figure 3.**
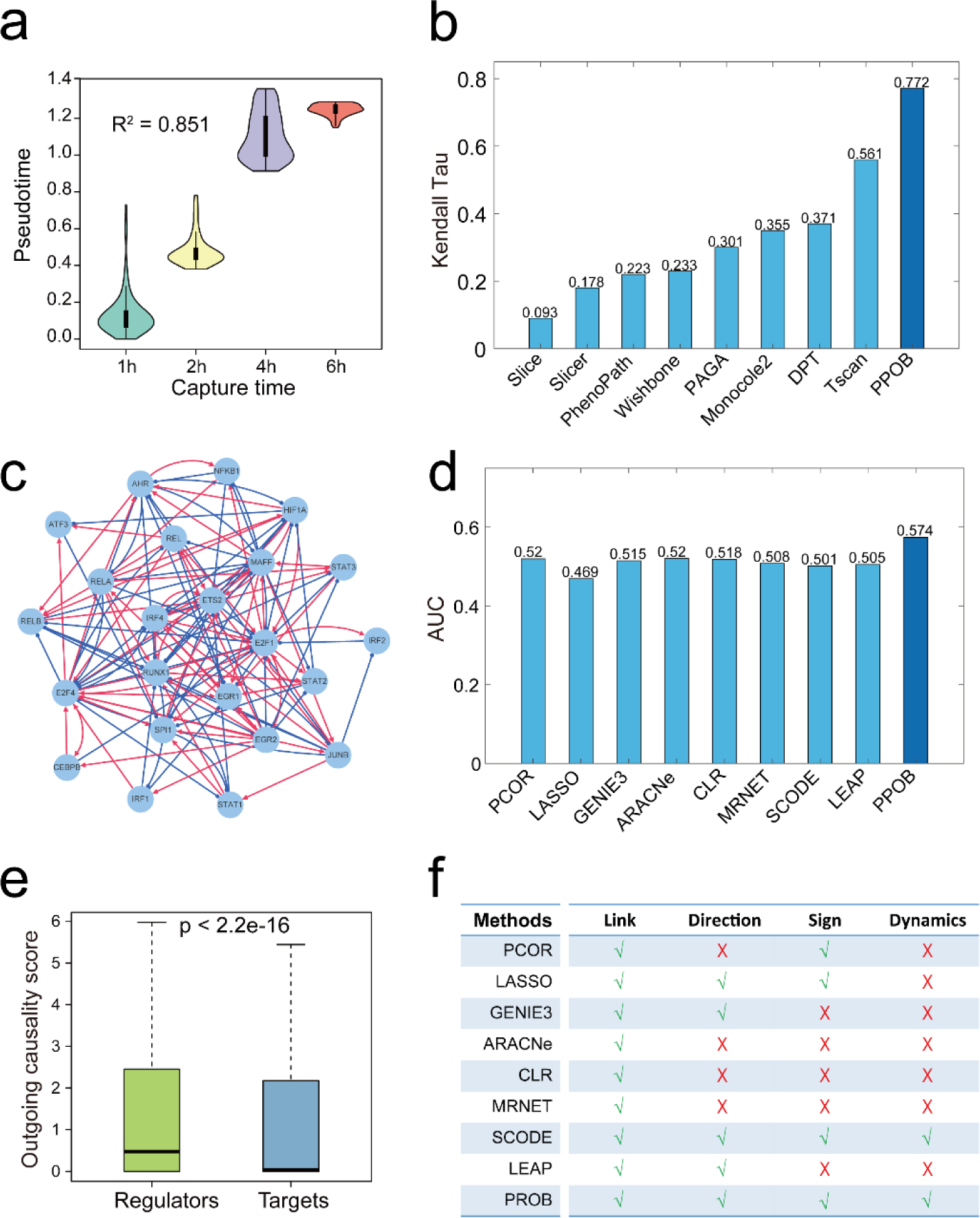
Comparison of PROB with other existing pseudotime inference methods and GRN inference methods using a real dataset. We employed a set of scRNA-seq data of dendritic cells (DCs) for benchmarking since the gold standard in this situation is available. The cells were sequenced at 1, 2, 4 and 6h after stimulation of LPS. (**a**) The estimated latent-temporal progression of cells recapitulated the real progression with R^2^ =0.851 to the capture times. (**b**) Benchmarking PROB with other pseudotime inference methods (Slice, Slicer, PhenoPath, Wishbone, PAGA, Monocole2, DPT, Tscan) evaluated by Kendall Tau and R^2^ (Fig S4). (**c**) a TF network inferred by PROB. (**d**) Benchmarking PROB with eight existing GRN inference methods (PCOR, LASSO, GENIE3, ARACNe, CLR, MRNET, SCODE and LEAP) based on an experimentally-defined TF network [43] evaluated by AUC of ROC. (**e**) PROB correctly revealed the ordering of the outgoing causality scores (on a log10 scale) for the known regulators and targets [38] on the DC scRNA-seq dataset. (**f**) Comparing properties of different methods in their capabilities of predicting network links, regulatory directions and signs as well as gene expression dynamics.

We next compared the accuracy of PROB with other existing GRN inference methods (e.g., PCOR, ARACNe, CLR, MRNET, Lasso, GENIE3, SCODE and LEAP) for cross-sectional data. A previous study measured binding region coverage scores for 23 TFs and thus quantified their regulatory potential in the DCs using a high-throughput Chromatin ImmunoPrecipitation (HT-ChIP) method [43]. A TF network was defined where an edge was viewed to be present if the coverage score between two TFs was greater than 0.3. We employed this network as a benchmark to compare the prediction accuracy of the network topologies inferred by PROB (**Figure 3c**) and other methods based on the above scRNA-seq data of DCs. The AUC values (**Figure 3d**) indicated that PROB outperformed the other existing methods.

Furthermore, we collected a set of known regulators and targets [38] to test whether PROB could correctly reveal the regulatory causality. To this end, we applied PROB to infer a GRN for 6 regulators and 28 targets based on the above DC scRNA-seq data and defined outgoing causality score (OCS) for each gene in the inferred network (see definition of OCS in the Methods section). The OCS values of regulators were much higher than that of targets (**Figure 3e**), suggesting that PROB faithfully revealed the ordering of the OCS values for the known regulators and targets on the analyzed dataset.

In addition, we summarized and compared the capabilities of the above methods in predicting gene regulatory links, directions, signs and expression dynamics (**Figure 3f**). Only PROB can simultaneously fulfill those four tasks in GRN inference.

### Reconstructing EMT regulatory networks during bladder cancer progression

Sarcomatoid urothelial bladder cancer (SARC) is a highly lethal variant of bladder cancer and has been reported to be evolved by the progression of the conventional urothelial carcinoma (UC) [39]. It has been demonstrated that the dysregulation of genes involved in the epithelial-to-mesenchymal transition (EMT) drives the progression of UC to SARC. To elucidate the dynamic change of the EMT regulatory network during the progression, here, we applied PROB to an expression dataset of bladder cancer containing 84 UC samples and 28 SARC samples (GSE128192). We collected 44 representative genes involved in several typical EMT-regulating pathways (**Table S1**). The expression patterns of these genes were recovered along with the inferred temporal progression (**Figure 4a**).

**Figure 4.**
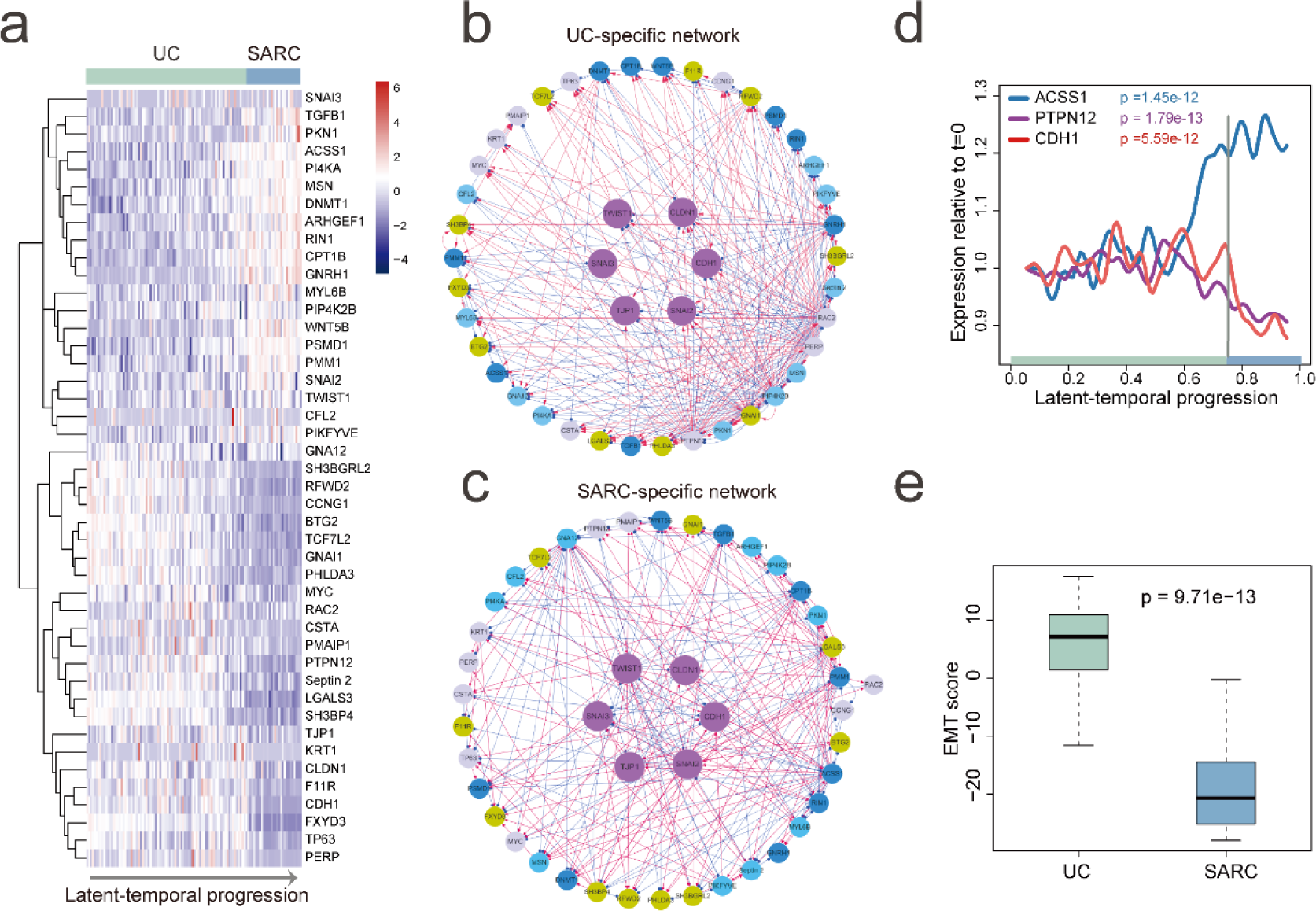
Reconstructing EMT regulatory networks during bladder cancer progression. (**a**) Expression patterns of the EMT regulatory genes along with the inferred latent-temporal progression of conventional urothelial carcinoma (UC) to aggressive sarcomatoid urothelial bladder cancer (SARC). (**b**) UC-specific network with edges unique to the UC network. (**c**) SARC-specific network with edges unique to the SARC network. Different colors of nodes in the network denote genes in different pathways (**Table S1**). (**d**) Reconstructed expression dynamics of ACSS1, PTPN12 and CDH1. ACSS1 and PTPN12 have largest out-degree values in the UC-specific network and SARC-specific network, respectively. CDH1 is a marker gene of epithelial state during EMT. (**e**) A decrease in EMT score indicated a transition from epithelial to mesenchymal state during the progression of UC to SARC. The EMT score for each tumor sample was calculated as weighted sum of expression levels of 73 EMT-signature genes as introduced in [39]. Positive EMT score corresponds to the epithelial phenotype while negative score to mesenchymal phenotype. Wilcoxon rank sum test (one-tailed) p value was calculated to assess the statistical significance.

We then applied PROB to reconstruct GRNs for UCs and SARCs, respectively, based on the ordered expression data of the above 44 genes. **Figure 4b** and **Figure 4c** show the UC-specific network and the SARC-specific network, respectively, suggesting rewiring of the EMT regulatory network during the progression of UC to SARC. The two networks were enriched with crosstalks between different pathways, indicating cooperative regulation of EMT by those pathways. PTPN12 and ACSS1 were found to have largest out-degree values in UC-specific network and SARC-specific network, respectively (**Table S1**). Temporal dynamics of gene expression (**Figure 4d**) showed that ACSS1 and PTPN12 oscillated synchronously with CDH1 (coding gene of epithelial marker protein E-cadherin) at the early stage of UC development. However, at a later stage before transition to SARC, ACSS1 dramatically increased and PTPN12 decreased. Meanwhile, the decrease of CDH1 later on indicated a transition from epithelial to mesenchymal phenotype in SRACs, in consistent with changes in EMT score values (**Figure 4e**).

### Validation of the role of ACSS1 in EMT

The decrease in PTPN12 expression during the progression is consistent with the previous finding that the loss of PTPN12 promotes EMT process and cell migration [44]. Furthermore, our result suggests that the up-regulation of ACSS1 might play a crucial role in the bladder cancer progression by promoting EMT program. We managed to validate the role of ACSS1 in EMT during bladder cancer progression, which has not been reported previously. The overexpression of ACSS1 in the 5637 cell line resulted in a significant decrease in CDH1 expression level (**Figure 5a**), and ACSS1 knockdown by small interfering RNA leaded to significant increase in CDH1 expression level (**Figure 5b**). The consistent changes in CDH1 protein levels following ACSS1 overexpression and knockdown were also observed (**Figure 5c-d**). These results confirmed that ACSS1 promoted EMT in bladder cancer cells. Furthermore, the immunohistochemical staining of patient samples (**Figure 5e**) revealed that conventional UC tumors showed focal retention of epithelial marker protein E-cadherin while SARC tumors showed focal retention of ACSS1, supporting the above estimated dynamics of ACSS1 and CDH1 during bladder cancer progression.

**Figure 5.**
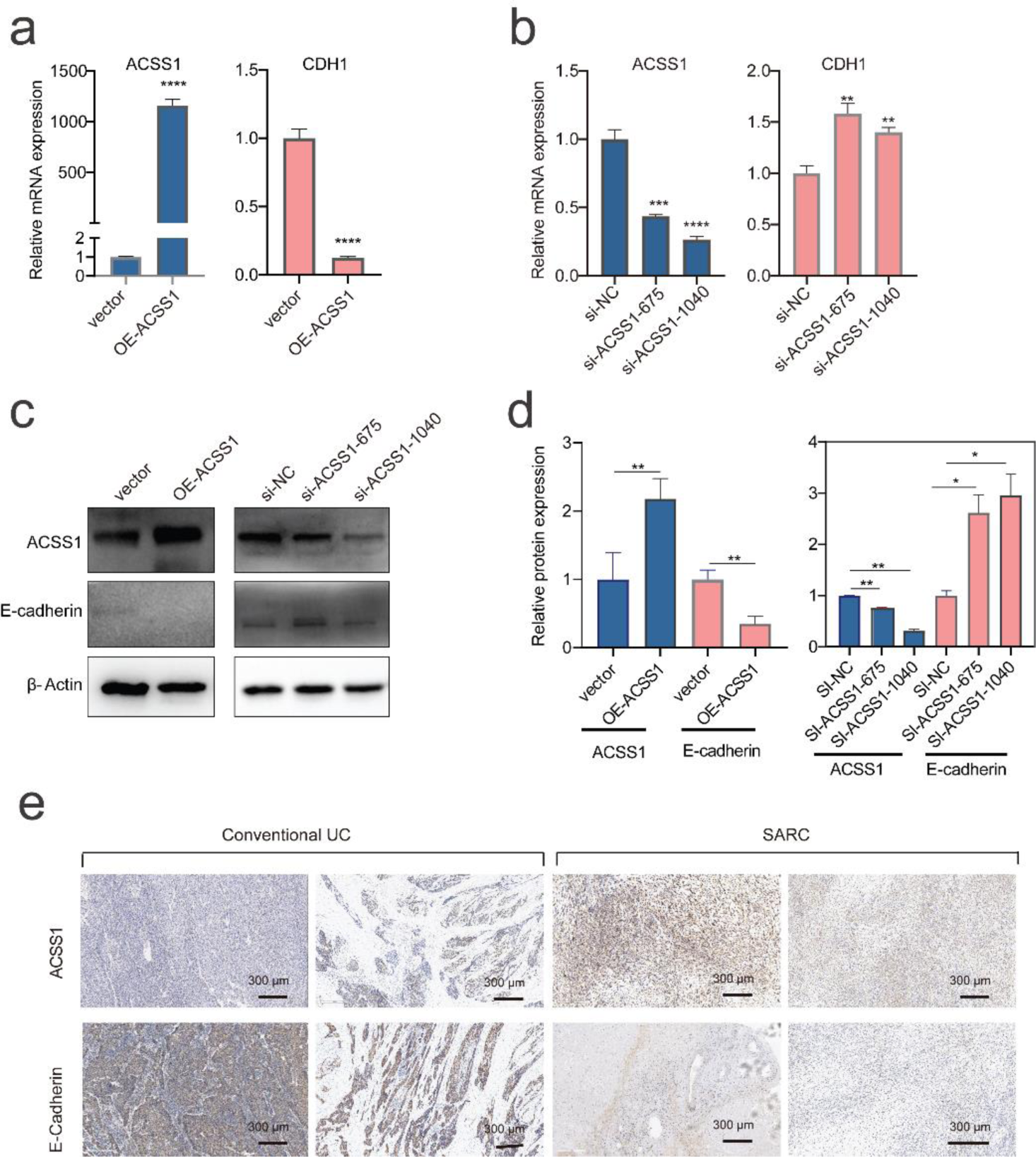
Experimental validation of the predicted role of ACSS1 in EMT of bladder cancer. (**a-b**) Expression levels of ACSS1 and CDH1 in 5637 cells when ACSS1 was overexpressed (a) and inhibited (b), measured by q-PCR. (**c**) Protein expression levels of ACSS1 and CDH1 in 5637 cells when ACSS1 was overexpressed or inhibited, measured by Western-blotting. (**d**) Quantification of the relative protein expressions. (**e**) Examples of immunohistochemical expression of ACSS1 and E-cadherin in conventional UC and SARC. Statistical significance was assessed by student’s t test. **P<0.01; ***P<0.001; ****P<0.0001. OE-ACSS1: overexpression of ACSS1; si-NC: small interfering RNA negative control; si-ACSS1: small interfering RNA targeting ACSS1.

### Identifying key gene regulators underlying breast cancer progression

To test whether our approach could be used to identify key genes underlying cancer progression, we applied PROB to a set of microarray data of breast cancer patients (n=196) with clinical information (GSE7390) (see details in **Text S5**) [40]. Based on the expression data reordered by PROB, we investigated which genes were upregulated or downregulated over progression by using a trend analysis technique. Such genes are referred to as temporally changing genes (TCGs) in this study. The one hundred top TCGs were selected. A heatmap with hierarchical clustering (**Figure 6a**) showed that these 100 genes were clearly clustered into two groups: a descending group (purple) and an ascending group (blue). We investigated the enriched gene sets for the two groups of genes using GSEA software [45, 46]. The descending genes were enriched in locomotion and movement of cell or subcellular component (**Figure 6b**, upper panel), and the ascending genes were mainly enriched in cell cycle and cell division processes (**Figure 6b**, lower panel).

**Figure 6.**
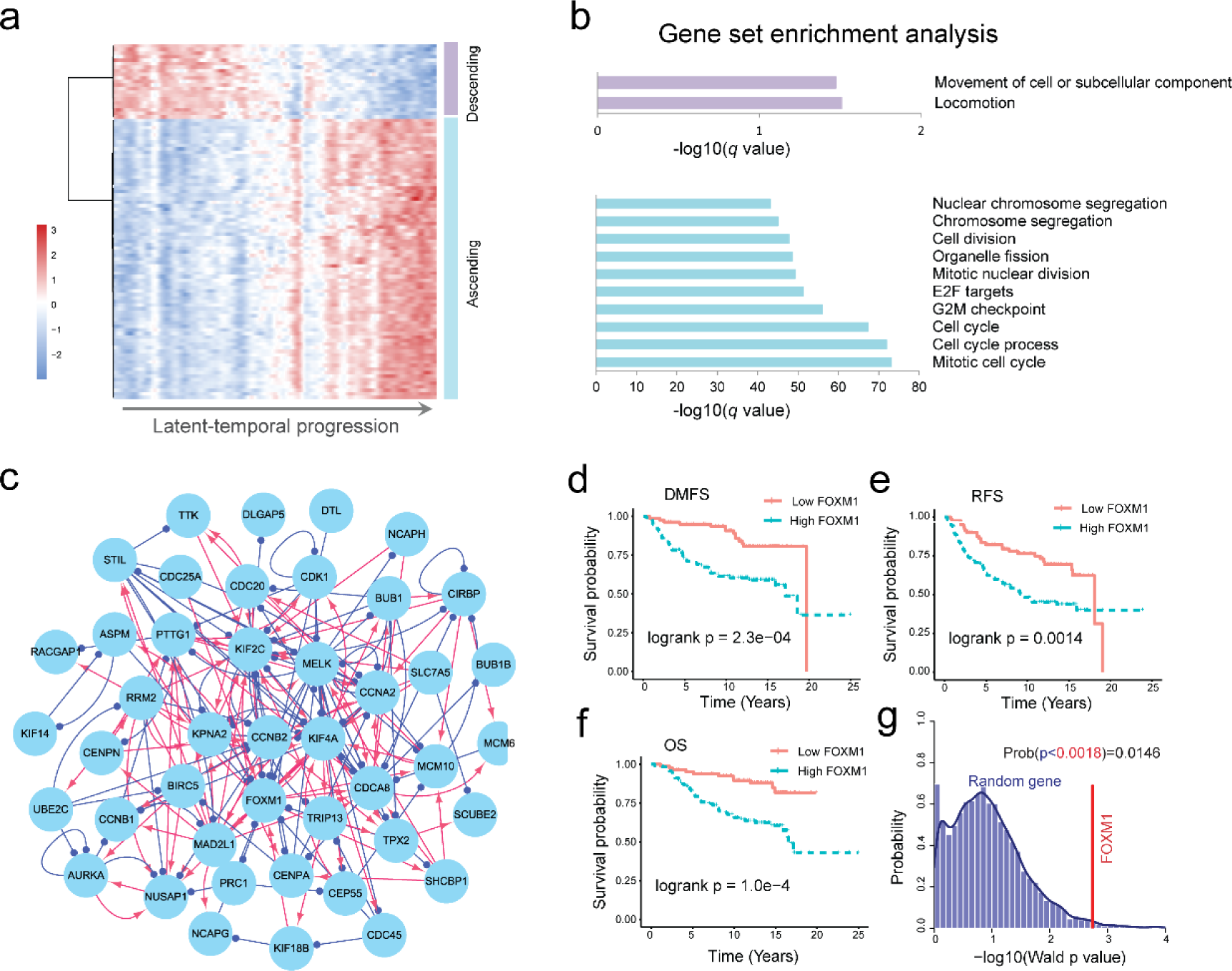
FOXM1 was revealed as a key gene underlying breast cancer progression by PROB. The gene expression data of 196 patients with clinical information (e.g., grade) were extracted from the GEO database (GSE7390 [40]). (**a**) Heatmap showing the expression profile of 100 selected genes that were most sustainably ascending (blue group) or descending (purple group) during cancer progression. (**b**) Gene set enrichment analysis for the descending genes (upper panel) and ascending genes (lower panel). The descending genes were enriched in local movement processes, and the ascending genes were mainly enriched in cell cycle and cell division processes. (**c**) The inferred GRN for the 100 genes. FOXM1 was found to be a hub gene in the network. (**d-f**) Clinical relevance of FOXM1 for breast cancer patients with respect to distant metastasis-free survival (DMFS) (**d**), relapse-free survival (RFS) (**e**) and overall survival (OS) (**f**). (**g**) Significance test of the prognostic power of FOXM1 using a bootstrapping approach. The *p* value from the permutation test was 0.0146, verifying the statistical significance of the prognostic power of FOXM1.

We then inferred the regulatory network of the above 100 top genes (**Figure 6c**). Based on an eigenvector centrality measure (**Text S5**), FOXM1 was identified as a most influential gene in the network. We found significant associations between FOXM1 and the distant metastasis-free survival (DMFS), relapse-free survival (RFS) and overall survival (OS) (**Figure 6d-g**) and therapeutic responses (**Figure S6**) in breast cancer patients (see details in **Text S6**). Previously, both *in vitro* and *in vivo* experiments have verified that FOXM1 plays important roles in promoting cell proliferation and cell cycle progression in breast cancer [47, 48]. Moreover, FOXM1 has been used as a key drug target in breast cancer [49, 50], and several drugs (e.g., daunorubicin, doxorubicin, epirubicin, and tamoxifen [51]) developed to target or inhibit FOXM1 have been tested in clinical trials (https://clinicaltrials.gov/). These evidences suggest that our network inference and analysis approach is effective to identify key genes of cancer progression or candidate drug targets.

### Validation of the FOXM1 subnetwork

A subnetwork was reconstructed for FOXM1, which predicted that FOXM1 could positively regulate ASPM, CDCA8, KIF2C, MCM10, MELK, NCAPG, SHCBP1 and STIL (**Figure 7a**). Preliminary investigation indicated that, except for STIL, the other 7 genes were functionally associated with FOXM1 according to String (https://string-db.org/), a database of functional protein-protein interaction networks (**Figure S7**). We proceeded to validate the expression changes of these predicted target genes using microarray data of MCF-7 cells that were treated with DMSO (control) or thiostrepton (a FOXM1 inhibitor) for 6 hours (GSE40766 [42]). We found that, except for SCCBP1 and STIL, the other 6 genes were significantly downregulated after FOXM1 inhibition (**Figure 7b**). The statistical significance was assessed using Wilcoxon rank-sum test (one-tailed) *p* values. These results suggest that PROB well predicted both the directions and signs of the edges in the FOXM1 subnetwork.

**Figure 7.**
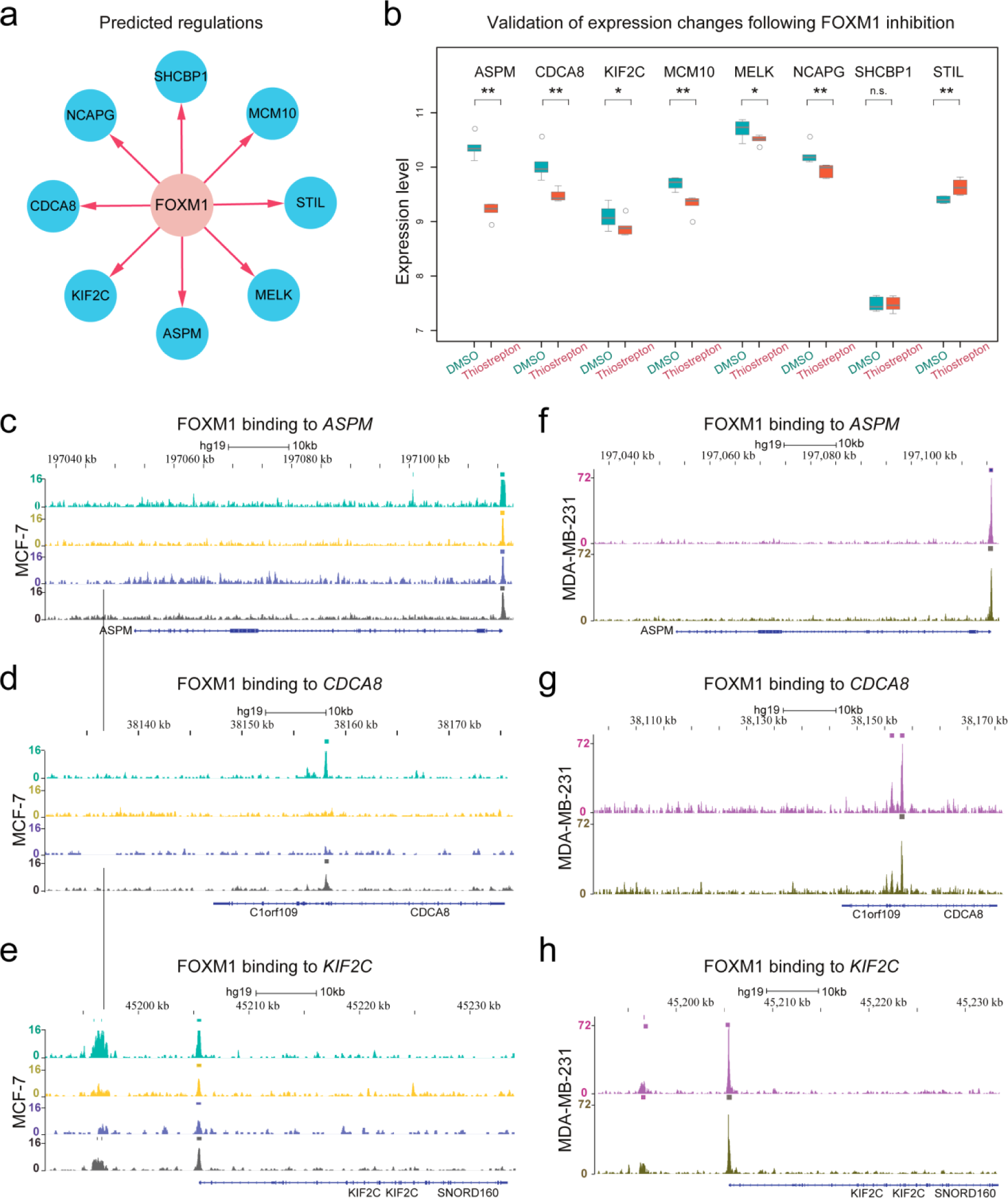
Validation of the predicted FOXM1 subnetwork. (**a**) The subnetwork of FOXM1 with predicted target genes. (**b**) Validation of the expression changes of the predicted target genes of FOXM1 with perturbation experiments. MCF-7 cells were treated with DMSO (control) or thiostrepton (a FOXM1 inhibitor) for 48 hours. Except for SCCBP1 and STIL, the other 6 genes were significantly down-regulated after FOXM1 inhibition. (**c-e**) ChIP-seq analysis of FOXM1 in the MCF-7 cell line with four biological replicates, showing that FOXM1 binds ASPM, CDCA8 and KIF2C. (**f-h**) ChIP-seq analysis of FOXM1 in the MDA-MB-231 cell line with two biological replicates, showing that FOXM1 binds ASPM, CDCA8 and KIF2C.

Moreover, we used ChIP-seq data (GSE40762 [42]) to analyze the binding of FOXM1 to the predicted targeted genes (**Text S7**). Both estrogen-dependent ER (+) MCF-7 and estrogen-independent ER (-) MDA-MB-231 human breast cancer cell lines were used for analysis. The analysis showed that FOXM1 binds ASPM, CDCA8 and KIF2C in both cell lines (**Figure 7c-h**). We note that the above three targets of FOXM1 were not previously reported by the widely used databases of transcriptional factor targets (e.g., TRANSFAC [52] and TRRUST v2 [53]). Interestingly, in another human mammary epithelial cell line (HMEC) (GSE62425 [54]) (**Figure S8**), the binding of FOXM1 to CDCA8 was absent, suggesting the emerging binding of FOXM1 to certain genes during the formation of breast cancer. In addition, we confirmed that the expression levels of the above three genes, ASPM, CDCA8 and KIF2C, were significantly reduced following the knockdown or silencing of FOXM1 based on both microarray data in BT-20 breast cancer cells (GSE2222 [55]) (**Figure S9a-c**) and RNA-seq data in MCF-7 breast cancer cells (GSE58626 [56]) (**Figure S9d-f**). These findings suggest that FOXM1 not only positively regulates the expression of but also directly binds to some of the predicted genes.

## Discussion

PROB provides a novel tool for inferring cancer progression and GRNs from cross-sectional data. Our approach is based on a dynamical systems representation of gene interactions during cancer progression. The inverse problem with respect to GRN reconstruction was solved by combining latent progression estimation and Bayesian inference for high-dimensional dynamic systems. PROB can be used to generate experimentally testable hypotheses on the molecular regulatory mechanisms of gene regulation during cancer progression and to identify network-based gene biomarkers for predicting cancer prognosis and treatment response.

Besides cross-sectional bulk transcriptomic data, our method can be naturally applied to time-course scRNA-seq data (**Figure 3**). Although scRNA-seq data can be used to infer GRNs during cell differentiation or development, it is currently not feasible to use scRNA-seq to investigate long term cancer progression due to patient heterogeneity, difficulty in acquisition of massive samples and expensive cost. In view of this, clinical transcriptomic data of cancer patients provide an alternative way to infer GRNs underlying cancer progression. The novelty and superiority of PROB can be first attributed to the successful ordering of tumor samples by using both gene expression data and staging information. Our proposed stage-weighted Gaussian kernel allows construction of diffusion-like random walks to quantify the temporal progression distance (TPD) between two patients (Equation (4)). The diffusion map, as a manifold-based nonlinear dimension reduction method, has been recently applied to scRNA-seq data analysis [25, 57, 58]. One major difficulty in applying diffusion maps for inferring pseudo trajectories lies in identifying the rooting point when using scRNA-seq data itself, and it often needs additional biological knowledge. An advantage of clinical transcriptomic data is that staging or grading information is usually available for samples as well, allowing development of an algorithm that automatically identifies the rooting point (Equation 5). We demonstrated that incorporating staging information into the temporal progression inference significantly improved its accuracy (**Figure S1**) and that our method significantly outperformed existing pseudotime inference methods (**Figure 3b** and **Figure S5**).

Considering technical variabilities in the sample-based transcriptomic data, it is important to have good robustness of the interaction coefficients in the GRN model with respect to the perturbation of the temporal progression. In addition to proving such property mathematically, through simulations we found PROB inference of both the progression trajectory and the gene network structure is rather robust to noise in the data (**Figure 2, Figure S4**). In addition, PROB is computationally efficient for GRN inference, which could be completed within 1 minute on the three real datasets analyzed in this study (**Table S4**).

For clinical applications, our method can be used to identify key genes for early detection of cancer progression and design of therapeutic targets. By recovering the temporal dynamics of gene expression in terms of the disease progression, PROB provides insights into exploiting kinetic features of functionally important genes that may be used as predictive biomarkers or drug targets. In the case study of bladder cancer progression, we have demonstrated that ACSS1 and PTNT12 played important roles in EMT during bladder cancer progression from UC to SARC and their expressions dynamically changed over the progression (**Figures 4-5**).

Therefore, we hypothesized that the temporal dynamics of EMT regulatory genes (e.g., ACSS1 or PTPN12) could be exploited to predict cancer progression. To this end, a logistic regression model was developed to predict EMT states or histological subtypes (UC vs. SARC) of bladder cancer based on the expression levels of ACSS1 and PTPN12, which showed good predictive accuracy (**Figure S10**). As such, the early changes in expressions of ACSS1 and PTPN12 during the progression of UC to SARC may be relevant for the early detection of SARC.

In another case study of breast cancer, FOXM1, a drugable target, was identified as a key regulator underlying breast cancer progression (**Figure 6**) and, importantly, the predicted FOXM1-target regulations were validated (**Figure 7**). Furthermore, here, we propose a GRN kinetic signature (**Text S8**) based on FOXM1-targeted gene interactions to prognosticate relapse in breast cancer. Kaplan-Meier (K-M) survival curves were plotted for the high-risk group (blue) and low-risk group (red) of patients with respect to relapse-free survival (RFS) (**Figure S11a-c**). The log-rank test *p* values for all three datasets were less than 1e-4. Moreover, we tested the statistical significance of the FOXM1-targets interactions in predicting relapse in breast cancer using a bootstrapping approach (**Text S8**). We compared the prognostic power (Wald test *p* value) of the FOXM1-predicted targets with that of 10000 sets of 8 randomly selected genes. The permutation test *p* values for all three datasets were less than 0.05 (**Figure S11d-f**), verifying the nonrandomness of the predicted targeted genes of FOXM1. These results demonstrated that the predicted FOXM1-target interactions could be used as a biomarker for prognosticating relapse in breast cancer. The latent-temporal progression–based casual network reconstruction method proposed in this study will likely innovate other network-based methodologies, such as those in system genetics [59, 60], network pharmacology [61, 62], and network medicine [1, 63].

Our method has several limitations that could be improved in future studies. For example, in the current method, only gene expression profiles and staging information from patient samples have been used for latent-temporal progression modeling. Other covariates, for example, age, genetic mutation, and molecular subtypes, might also be useful for progression inference [64]. Statistical models that integrate multiple aspects of clinical information will provide better inference of disease progression.

In summary, we have developed a novel latent-temporal progression-based Bayesian Lasso method, PROB, to infer directed and signed gene networks from prevalent cross-sectional transcriptomic data. PROB provides a dynamic and systems perspective for characterizing and understanding cancer progression based on patients’ data. Our study also sheds light on facilitating the regulatory network-based approach to identifying key genes or therapeutic targets for the prognosis or treatment of cancers.

## Acknowledgements

We would like to acknowledge Prof. Tianshou Zhou at Sun Yat-sen University for valuable discussion. We would also like to acknowledge Dr. Zifeng Wang at Sun Yat-sen University Cancer Center for providing information on the Chip-seq data and Dr. Dongliang Leng at the University of Macau for processing the Chip-seq data.

## Data availability

The gene expression dataset of the bladder cancer was downloaded from the NCBI GEO database (GSE128192). The gene expression datasets as well as clinical information of the breast cancer patients used for network prediction were downloaded from the NCBI GEO database (GSE7390). The microarray and ChIP-seq data used for network validation were downloaded from the NCBI GEO database (GSE40766, GSE40762, GSE62425, GSE2222, GSE58626 and GSE27830). The clinical gene expression data used for survival analysis were downloaded from the NCBI GEO database (GSE2990, GSE12093, GSE5327, GSE1456, GSE2034, GSE3494, GSE6532 and GSE9195). Other datasets used for validating and benchmarking PROB were downloaded from the TCGA database (TCGA COAD and TCGA SKCM). A recent re-quantification of the LPS scRNA-seq dataset (GSE48968) was downloaded from the conquer database (http://imlspenticton.uzh.ch:3838/conquer/). The code for PROB is available at https://github.com/SunXQlab/PROB.

## Funding

XS was supported by grants from the National Natural Science Foundation of China (11871070), the Guangdong Basic and Applied Basic Research Foundation (2020B151502120), the Fundamental Research Funds for the Central Universities (20ykzd20) and Guangdong Key Field R&D Plan (2019B020228001). QN was partially supported by a National Science Foundation grant DMS1736272, a Simons Foundation grant (594598), and National Institute of Health grants U01AR073159 and U54CA217378.

## Supporting Information Legends

Supplementary information is available online, including the following contents:

**Text S1**. Method details of PROB.

**Text S2**. Proof of the Theorem 1.

**Text S3**. Implementation of PROB.

**Text S4**. Simulation study.

**Text S5**. PROB applied to realistic datasets.

**Text S6**. Clinical relevance of FOXM1 to breast cancer.

**Text S7**. Validation of the predicted FOXM1-targets interactions.

**Text S8**. GRN kinetic signature.

**Figure S1**. Incorporation of staging information significantly improved the accuracy of latent-temporal progression inference.

**Figure S2**. Validation of PROB against the synthetic datasets with different levels of variabilities.

**Figure S3**. Posterior distribution of regulatory parameters associated with Fig S1.

**Figure S4**. Evaluation indexes for inference of latent-temporal progression and GRN inference under different variability levels in the synthetic datasets.

**Figure S5**. Benchmarking PROB with other existing pseudotime inference methods.

**Figure S6**. FOXM1 expression was associated with the therapeutic responses of breast cancer patients.

**Figure S7**. The functional interaction network of FOXM1 extracted from String database.

**Figure S8**. ChIP-seq analysis of FOXM1 in human mammary epithelial cells (HMEC).

**Figure S9**. Another validation of the predicted regulation of KIF2C, ASPM and CDCA8 by FOXM1.

**Figure S10**. ACSS1 and PTPN12 are predictive of EMT and progression of UC to SARC.

**Figure S11**. A GRN kinetic signature predicts relapse in breast cancer.

**Table S1**. Out-degree values of the genes in the UC-specific and SARC-specific networks.

**Table S2**. The siRNA sequence used in this study.

**Table S3**. The specific primers used in this study.

**Table S4**. Runtime of PROB on different datasets.

**Supplementary References**

